# Identification of two cancer stem cell-like populations in triple-negative breast cancer xenografts

**DOI:** 10.1101/2021.10.21.465257

**Authors:** Jun Nakayama, Hiroko Matsunaga, Koji Arikawa, Takuya Yoda, Masahito Hosokawa, Haruko Takeyama, Yusuke Yamamoto, Kentaro Semba

## Abstract

Gene expression analysis at the single-cell level by next generation sequencing has revealed the existence of clonal dissemination and microheterogeneity in cancer metastasis. The current spatial analysis technologies can elucidate the heterogeneity of cell–cell interactions *in situ*. To reveal the regional and expressional heterogeneity in primary tumors and metastases, we performed transcriptomic analysis of microtissues dissected from a triple-negative breast cancer (TNBC) cell line MDA-MB-231 xenograft model with our automated tissue microdissection punching technology. This multiple-microtissue transcriptome analysis revealed three cancer cell-type clusters in the primary tumor and axillary lymph node metastasis, two of which were cancer stem cell (CSC)-like clusters (CD44/MYC-high, HMGA1-high). Reanalysis of public single-cell RNA-seq (scRNA-seq) datasets confirmed that the two CSC-like populations existed both in TNBC xenograft models and TNBC patients. In addition, the gene signature of the HMGA1-high CSC-like cluster has the potential to serve as a novel biomarker for diagnosis. The diversity of these multiple CSC-like populations may cause differential anticancer drug resistance, increasing the difficulty of curing this cancer.

## Introduction

Breast cancer cells metastasize to multiple distant organs, such as the axillary lymph nodes, lungs, bone, liver, and brain [1, 2]. In particular, metastasis to axillary lymph nodes is an indicator of cancer grade in breast cancer patients [3]. Most breast cancer tissues, including distant metastases, exhibit genetic heterogeneity [4]. Single-cell analyses have revealed that cancer cells evolve through the acquisition of genomic mutations in the primary tumor and metastases [5, 6]. Most previous analyses have been performed using isolated cancer cells and stromal cells from cancer tissues. Thus, the cell–cell interactions between cancer cells and stromal cells remain to be analyzed. In recent studies, current single-cell analysis and spatial transcriptome technologies have revealed the heterogeneity of cell–cell interactions between cancer cells and stromal cells *in situ* [7–9]; however, further analysis is needed to elucidate the nature of tumor heterogeneity.

Comprehensive gene expression analysis of metastases harvested from approximately 500 specimens of various cancer types and metastatic organs (MET500 cohort) has suggested that metastatic tissues can be divided into several categories (e.g. proliferative or EMT-like/inflammatory) [10]. In particular, some samples were found to show signatures of more than one category, suggesting that these samples have micro-intratumor heterogeneity. Heterogenous tumors contain a small subpopulation of cancer stem cells (CSCs) able to induce anticancer drug resistance and metastasis [11–13]. To clarify such heterogeneity, microtissue sectioning using laser capture microdissection has often been performed [14]. This method has several disadvantages, including the laborious and time-consuming nature of sample handling and a high risk of RNA degradation. Thus, in previous work, we developed a system involving automated tissue microdissection punching followed by transcriptomic analysis of the tumor microtissue [15]. This site-specific rapid sampling method by a hollow punching needle from frozen tissue enables to be performed on low-cost molecular analyses at low-resolution spatially resolved tissue specimens. To analyze the expressional heterogeneity in microtissues from the primary tumor and axillary lymph node metastasis, we performed analysis of the spatial microtissue transcriptome in a xenograft model with the triple-negative breast cancer (TNBC) cell line MDA-MB-231. We focused on the expression profiles of known metastasis-promoting genes and CSC markers in dissected microtissues.

## Results

### Sampling microtissues from primary tumor and axillary lymph node metastasis in MDA-MB-231 xenografts

Primary tumor and axillary lymph node metastasis were harvested from NOD-SCID mice with MDA-MB-231-parent-*Venus* cell line xenografts. We subjected the sliced tissues to microtissue dissection by an automated tissue microdissection punching system (Figure 1A). RNA was successfully recovered from the microtissues collected at 93 spots in the primary lesion and 44 spots in axillary lymph node metastasis using a microtissue automatic sampling device (Figure 1B). In samples of this size, although the number of cells present in the tumor tissue varies, it can be inferred that several to approximately 10-30 cells are present in each spot [15]. RNA-seq analysis was performed on the total RNA extracted from each spot. We checked the quality of the FASTQ files by FASTQC. Total RNA samples contained RNA from human cancer cell lines and RNA from mouse stromal cells in the tumor microenvironment. Therefore, the obtained sequences were mapped to both the human reference genome and the mouse reference genome by HISAT2 [16]. Protein-coding genes (human: 19961 genes, mouse: 22050 genes) were extracted as transcripts per million (TPM) for spatial transcriptome analysis with Seurat (Figure 1C, Supplementary Figure S1A, S1B and S1C) [17, 18].

**Figure 1.**
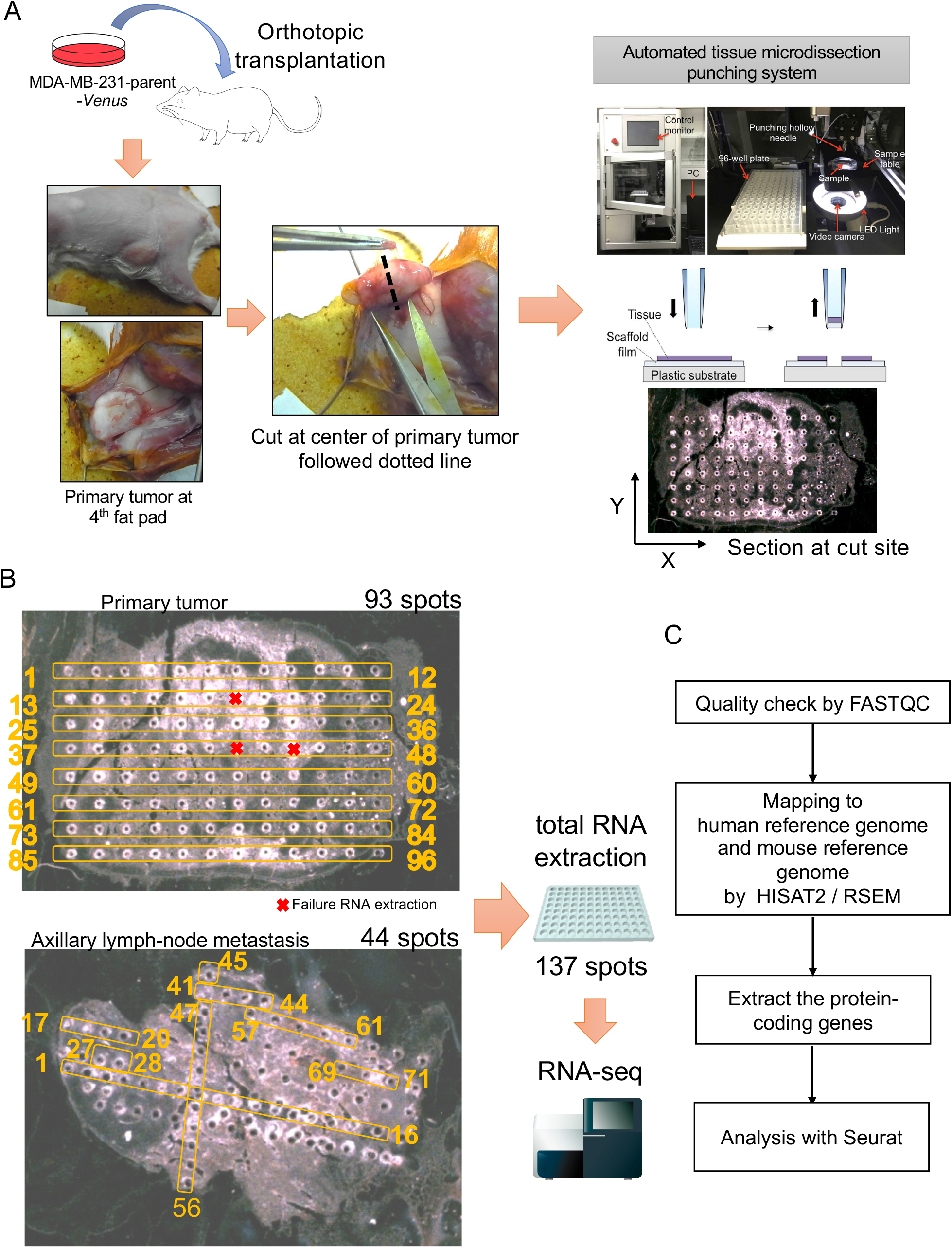
Microtissue sectioning of the primary tumor and the axillary lymph node metastasis from the TNBC xenograft model. (A) Experimental flowchart of the spatial transcriptomics analysis. The human triple-negative breast cancer (TNBC) cell line MDA-MB-231-Parent-Venus was transplanted orthotopically into a female NOD-SCID mouse. After 8 weeks, the primary tumor was harvested, and the center of the tumor was sectioned. After 4 weeks, axially lymph node metastases were harvested and sectioned from the same mouse. Sectioning was performed by an automated tissue microdissection punching system with a 100 μm needle. (B) In total, 93 microspots were sectioned from the primary tumor, and 43 microspots were sectioned from axillary lymph node metastases. RNA was extracted from a total of 137 spots. (C) Flowchart of the transcriptome analysis. A quality check was performed by FASTQC. The reads were mapped to the human reference genome and the mouse reference genome by HISAT2 and RSEM. Protein-coding genes were selected for analysis with Seurat.

### Microtissue transcriptomics analysis reveals two types of cancer stem cell-like populations

The clustering analysis and UMAP plots showed 3 clusters of cancer cells (transcripts mapped to the human reference genome) and 4 clusters of stromal cells (transcripts mapped to the mouse reference genome) in the microspots dissected from primary tumors and axillary lymph node metastases (Figure 2A, 2B and Supplementary Table S1). Next, we evaluated the expression of CSC markers to focus on the CSC populations in the primary tumor and metastatic lesion [13]. We found that human cancer clusters showed specific gene expression patterns for high mobility group AT-Hook1 (HMGA1) and CD44 (Figure 2C, 2E and Supplementary Table S2). CD44 and HMGA1 are well-known markers of CSCs in breast cancer [19, 20]. CD44 was broadly expressed in all human cell clusters; on the other hand, HMGA1 was highly expressed in only HMGA1-high clusters (Figure 2C). The upstream enrichment analysis using Metascape [21] showed that the differentially expressed genes (DEGs) of the CD44-expressing clusters were enriched in MYC-targeted genes (Supplementary Table S3). Consequently, these human cell clusters were named HMGA1-high, CD44/MYC-high, and marker-low. The DEGs in the human cell clusters were extracted and visualized as a heatmap (Figure 2D and Supplementary Table S2). The heatmap showed that the HMGA1-high cluster expressed TMSB10 [22], CTSD [23], and LGALS1 [24, 25], which are correlated with poor prognosis in breast cancer. The human CD44/MYC-high cluster expressed CENPK [26] and CENPN [27] which regulate the cell cycle and cell division in cancer. The human marker-low clusters showed low expression levels of these genes. These results suggested that two types of CSC-like populations existed in the MDA-MB-231 xenografts.

**Figure 2.**
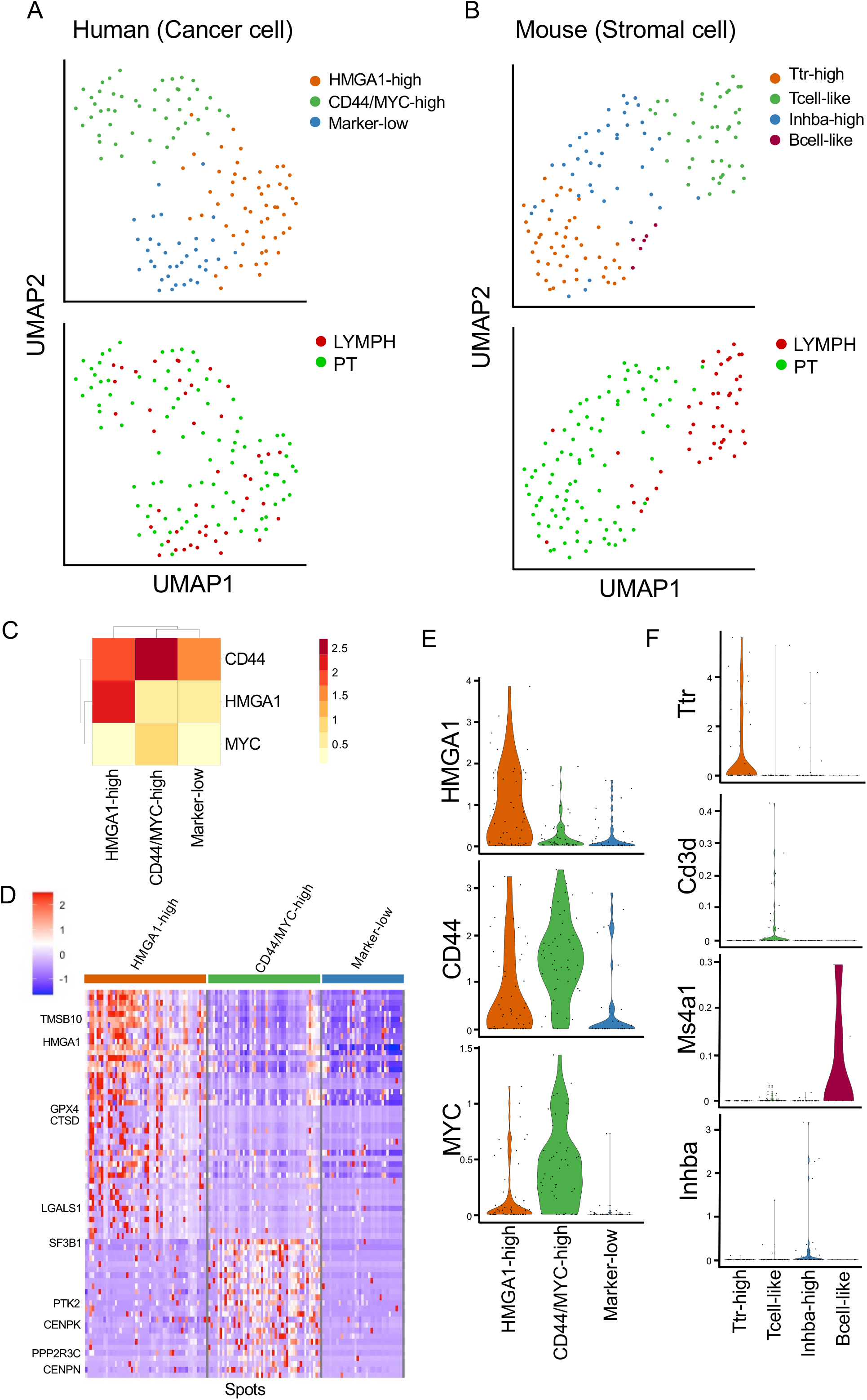
Transcriptome profiling of the microspot clusters. (A) UMAP plot of human (cancer cell) spot clustering. (B) UMAP plot of mouse (stromal cell) spot clustering. (C) Heatmap of CD44, HMGA1, and MYC expression in each human cell cluster. (D) Heatmap of differentially expressed genes (DEGs, adjusted p value < 0.05 and pct.1 - pct.2 > 0.1) in each human cell cluster. (E) Violin plots of cancer marker genes. (F) Violin plots of stromal marker genes.

To determine the biological function of DEGs, we performed upstream analysis, GO enrichment analysis, and pathway enrichment analysis using Metascape. We performed pathway and GO enrichment analyses focused on the two CSC-like clusters that had upregulated DEGs (Supplementary Figure S2). Amide metabolites, VEGFA-VEGFR signaling, and ribonucleoprotein complex biogenesis were enriched in both clusters. On the other hand, the DEGs in the CD44/MYC-high clusters were enriched in many terms related to the cell cycle, cell division and ribosomal biogenesis. In the HMGA1-high clusters, the terms ribosome and TRBP complex were significantly enriched.

The expression of CD44 and HMGA1 in transplanted sections was confirmed with fluorescence immunostaining (Figure 3A). CD44 and HMGA1 were detected in primary tumor sections. There were cells that expressed CD44 and HMGA1 independently and cells that coexpressed both CD44 and HMGA1. To confirm the existence of two CSC-like populations in the MDA-MB-231 xenograft model, we reanalyzed the public single-cell RNA-seq (scRNA-seq) samples of primary tumors and circulating tumor cells (CTCs) from the MDA-MB-231-LM2 xenograft model [28]. We downloaded the GSE163210 datasets from the Gene Expression Omnibus (GEO) database and reanalyzed it with Seurat/R (Figure 3B). We extracted 8,494 cancer cells from the dataset with UMAP visualization (Figure 3C). The scRNA-seq analysis revealed that there were HMGA1-high, CD44-high, and double-positive (HMGA1- and CD44-positive) populations in both the primary tumors and CTCs in the xenograft model (Figure 3D and 3E). These results suggested that there were two types of CSC-like populations expressing CSC marker genes in the MDA-MB-231 xenograft model.

**Figure 3.**
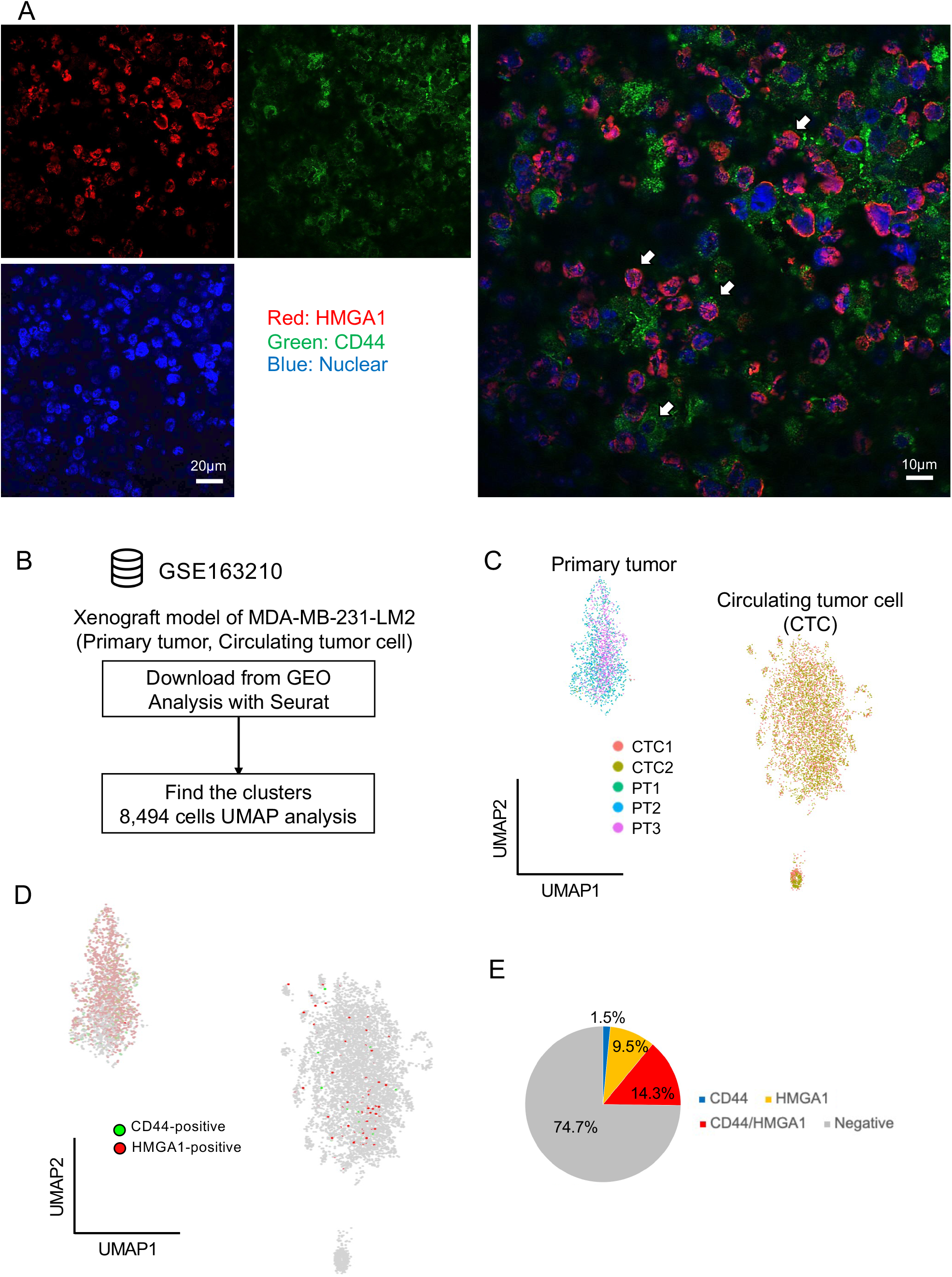
Two cancer stem cell-like populations detected by Immunostaining and scRNA-seq analysis. (A) Representative images of CD44 and HMGA1 immunostaining in primary tumor sections from the MDA-MB-231 xenograft model. Red: HMGA1, green: CD44, and blue: nucleus. Scale bars, 20 μm (left), 10 μm (right). White arrows represent cancer cells with independent expression or coexpression. (B) Flowchart of public single-cell RNA-seq (scRNA-seq) reanalysis using the GSE163210 dataset. (C) UMAP plot of MDA-MB-231-LM2 xenograft tumors and circulating tumor cells. (D) Expression of CD44 and HMGA1 in the UMAP plot. (E) Pie chart of the cancer stem cell (CSC)-like population proportion in MDA-MB-231-LM2 xenografts.

On the other hand, the mouse stromal clusters showed specific gene expression patterns for transthyretin (Ttr), Cd3d (T-cell marker), membrane-spanning 4-domains a1 (Ms4a1, B-cell marker), and inhibin subunit beta A (Inhba; a subunit of both activin and inhibin) (Figure 2F and Supplementary Figure S3A). Ttr and Inhba were highly expressed in their respective specific clusters; therefore, these mouse clusters were named Ttr-high, Tcell-like, Inhba-high, and Bcell-like. Enrichment analysis of the DEGs of the mouse clusters showed that only the Tcell-like clusters had many enriched terms (Supplementary Figure S3B and Supplementary Table S4). The citric acid (TCA) cycle, chemical stress response, and fatty acid oxidation were enriched in the Tcell-like populations.

The heterogeneity of each cluster was demonstrated by calculation of network topology using the normalized closeness centrality [29, 30]. The centralities showed the correlational cellular diversity based on gene expression in each cluster. These results showed that the cancer cells in the HMGA1 clusters had a high similarity of expressional diversity compared with those in the CD44/MYC clusters (Supplementary Figure S4A and S4B).

### Microspot spatial and cell cycle analyses

Next, we performed spot analysis with the spatial information to determine the spatial heterogeneity in the xenografts (Figure 4A, 4B, and 4C). Interestingly, although 3 human cell clusters were present in both the primary tumors and the lymph node metastases (Supplementary Figure S5A), the mouse stromal clusters showed a site-specific pattern (Supplementary Figure S5B). Most of the Ttr-high clusters were observed in the primary tumors. On the other hand, most of the Tcell-like clusters and Bcell-like clusters were found in the lymph node metastases. Human CD44/MYC-high cancer cells tended to localize the outside of the primary tumor (Figure 4A and 4D).

**Figure 4.**
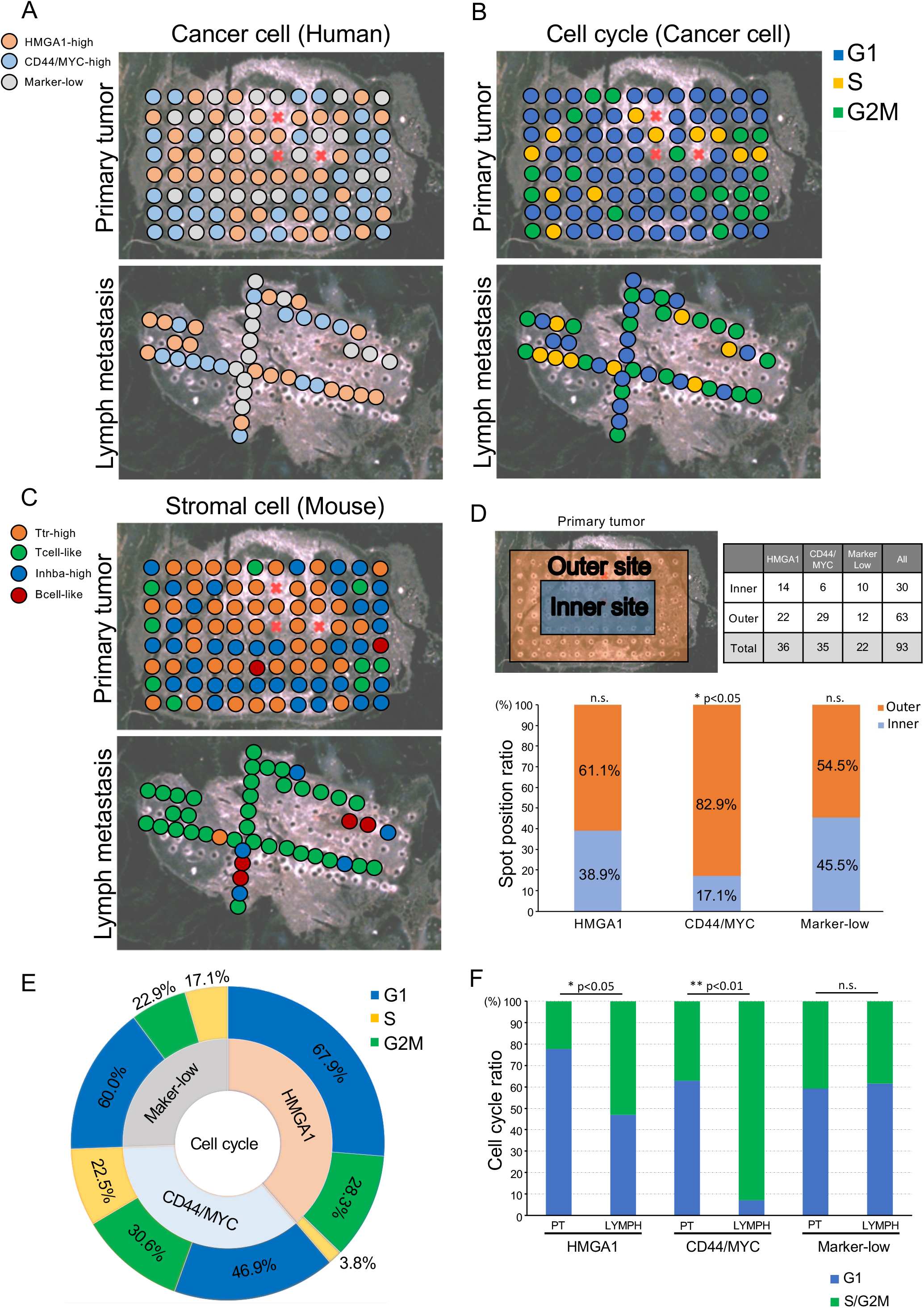
Spatial analysis of the two CSC-like populations. (A) Spatial transcriptomics analysis of human (cancer cell) clusters in the primary tumor and the lymph node metastasis. (B) Cell cycle phase of cancer cells in the primary tumor and lymph node metastases. (C) Spatial transcriptomics analysis of mouse (stromal cell) clusters in the primary tumor and lymph node metastasis. (D) Spatial analysis of human (cancer cell) clusters with the chi-test; *p < 0.05; n.s., not significant. (E) Sunburst plot of the cell cycle in human cancer cell clusters. (F) Bar plot of cell cycle phases in the primary tumor and lymph node metastasis. Fisher’s exact test, *p < 0.05; **p < 0.01; n.s., not significant.

Cell cycle analysis of the cancer cells showed that cell proliferation occurred outside of the primary tumor and at sparse sites among the lymph node metastases (Figure 4B). Approximately 50% of the cells in the CD44/MYC-high clusters and 30% of the cells in the HMGA1-high clusters were actively undergoing cell division (Figure 4E, Supplementary Figure S5C and Supplementary Table S5). These results suggested that the cells that had divided originated from the two CSC-like clusters, leading to cancer expressional heterogeneity. Next, a comparative analysis of the cell cycle in the primary tumor and lymph node metastasis showed that the two CSC-like clusters (HMGA1-high and CD44/MYC-high) had an increased cell division index in lymph node metastasis (Figure 4F). In contrast, the marker-low clusters did not have an altered cell cycle index in either location. These results suggested that the CD44/MYC and HMGA1 CSC-like populations in metastatic tissues proliferated aggressively.

Mouse stromal cell localization assessment showed that most of the Tcell-like clusters were present throughout the entire lymph node metastases; on the other hand, the Tcell-like clusters also existed outside the primary tumor. Most of the Ttr-high clusters were sparsely present throughout the primary tumor (Figure 4C). The cells in the mouse Tcell-like clusters and Inhba-high clusters showed an active cell cycle (Supplementary Figure S5D, S5E and S5F).

### Two cancer stem cell-like populations in TNBC patients

To confirm our findings in clinical samples, we reanalyzed the integrated 3 public scRNA-seq datasets [8, 31, 32] and data from the Molecular Taxonomy of Breast Cancer International Consortium (METABRIC) [33]. First, we analyzed the integrated scRNA-seq dataset of 19 TNBC samples (Figure 5A). We extracted 48,362 cancer cells from the dataset with UMAP visualization (Figure 5B and Supplementary Figure S6A). Expression analysis detected HMGA1-high cancer cells (HMGA1 expression level > 2), CD44-high cancer cells (CD44 expression level > 2), and double-positive cancer cells in the integrated TNBC cohorts (Figure 5C, 5D, and Supplementary Figure S6B). Most of the TNBC patients had two CSC-like populations and a double-positive population (Figure 5E, 5F and Supplementary Table S6).

**Figure 5.**
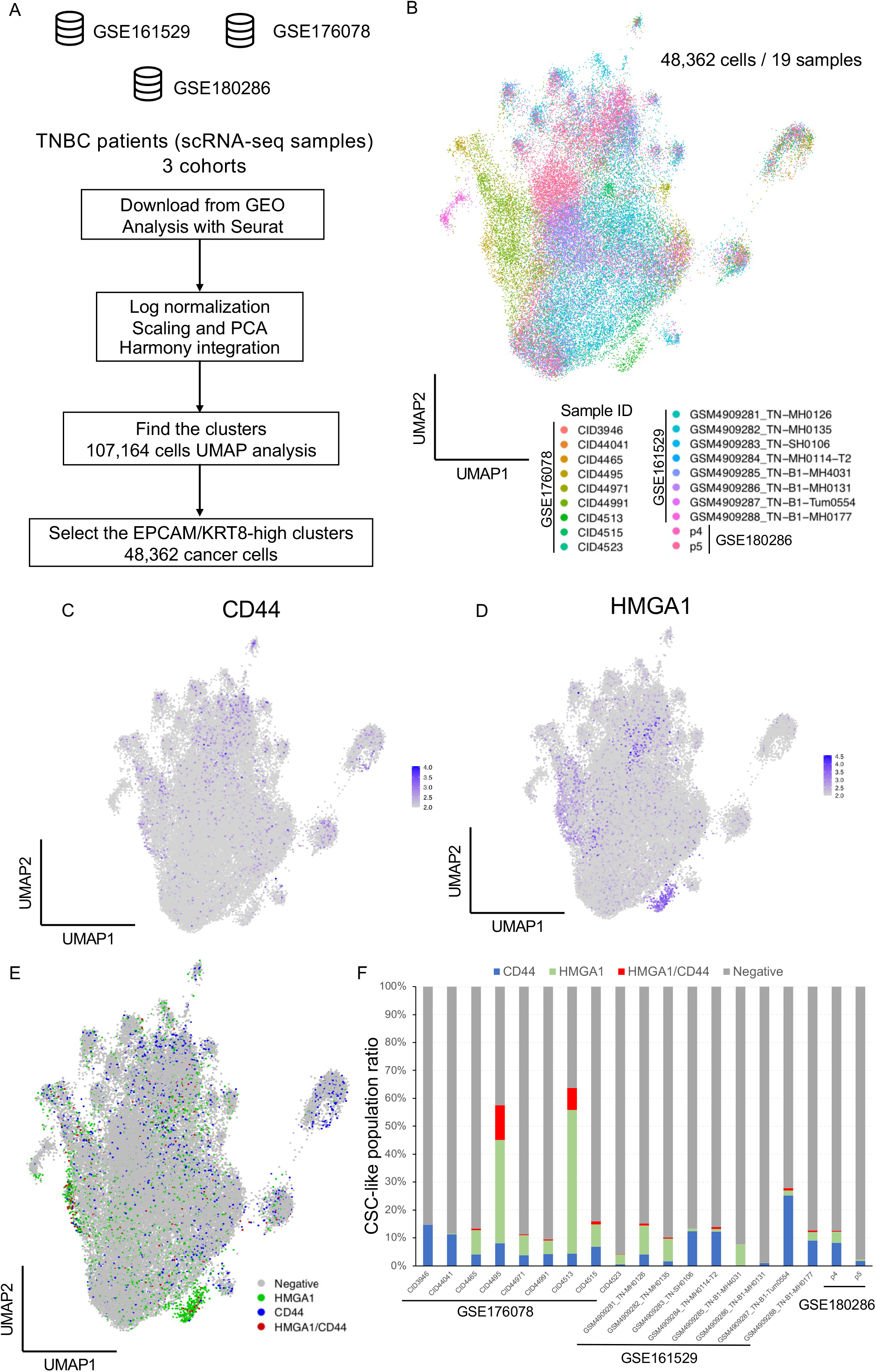
Reanalysis of clinical scRNA-seq with CSC-like signatures. (A) Flowchart of the reanalysis of a public scRNA-seq datasets. We downloaded GSE161529, GSE176078, and GSE180286 (scRNA-seq data from 19 TNBC patients). The integrated datasets were analyzed with Seurat. Log normalization, scaling, PCA and UMAP visualization were performed following the basic protocol in Seurat. To extract the cancer cells, cells expressing EPCAM/KRT8 (epithelial marker) were filtered. (B) UMAP plot of cancer cells from 19 TNBC patients. (C&D) Expression analysis of CD44 (expression level > 2) and HMGA1 (expression level > 2) with UMAP plots. (E) UMAP plot of CD44-high, HMGA1-high, HMGA1/CD44-high, and marker-negative cancer cells. (F) The bar plot shows the proportion of cancer cells that expressed CD44 and HMGA1.

Next, survival analysis was performed using the HMGA1-high gene signatures and CD44/MYC-high gene signatures with METABRIC TNBC cohorts (398 patients). The HMGA1 gene signatures and CD44/MYC gene signatures were selected from the DEGs analysis (adjusted p value < 0.05) excluding ribosomal genes (Supplementary Table S7). High expression of HMGA1 signatures correlated with poor prognosis in the claudin-low subtype (Figure 6A). In contrast, CD44/MYC signatures did not correlate with prognosis in these cohorts (Figure 6B).

**Figure 6.**
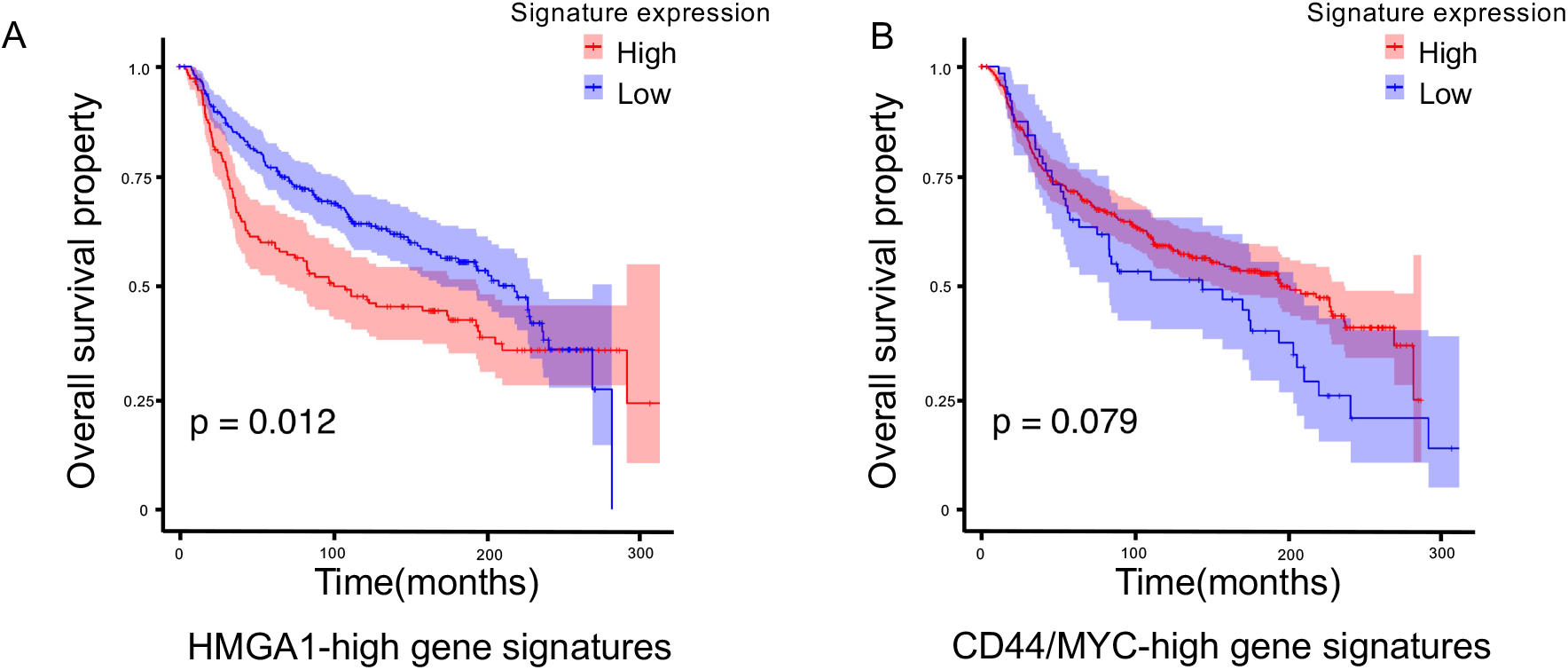
Survival analysis based on the HMGA1-high and CD44/MYC-high expression gene signatures. (A) Survival analysis of TNBC patients (claudin-low subtype and basal-like subtype) in METABRIC cohorts with the Kaplan–Meier method. Survival analysis according to the expression of HMGA1 signatures (high = 151, low = 247). Shading along the curve indicates the 95% confidential interval. Log-rank test, p = 0.012. (B) Survival analysis based on the expression of CD44/MYC signatures (high = 333, low = 65). Log-rank test, p = 0.079.

### Cell–cell interaction (CCI) analysis of microspots

CSC and stromal niche cell interactions are important events related to drug resistance and metastasis [11, 34]. A method to estimate the cell–cell interactions in each spot (detailed in the materials and methods) was developed for the evaluation of cell–cell interactions in the microtissues of the primary tumor and metastatic lesions. This analysis will be helpful for assessing the CSC niche and metastatic niche [13]. To estimate the CCI score, we utilized a cell–cell interaction database (https://baderlab.org/) and extracted a total of 115,900 interactions (3,209 ligands, 4,364 receptors and 433 extracellular matrix components) [29, 30]. Highly expressed genes (expression level > 2) were selected for CCI analysis, and ligand–receptor interactions (human to mouse and mouse to human) were estimated in each spot (Supplementary Figure S7A). We extracted 2,432 interactions and 7 clusters of “mouse (stromal) to human (cancer)” CCI patterns (Supplementary Figure S7B and Supplementary Table S8). In particular, Cluster 2 was present in only lymph node metastasis spots, while Cluster 6 was present in by only primary tumor spots. We thus focused on these clusters: the CCI heatmap showed lymph node metastasis-specific CCIs (Supplementary Figure S7C and S7D) and primary tumor-specific CCIs (Supplementary Figure S7E and S7F). The “human (cancer) to mouse (stromal)” CCI pattern showed 741 interactions (Supplementary Figure S8 and Supplementary Table S8).

## Discussion

Spatial transcriptomics technologies have enabled us to reveal the *in situ* expressional profiles and microheterogeneity of cancer [9]. In particular, in a xenograft model, both human-derived RNA and mouse-derived RNA can be analyzed both simultaneously and individually by mapping the sequence reads to a human genome reference or mouse genome reference [35, 36]. In this study, by combining microtissue sampling and the isolation of human-mouse gene expression by mapping, we revealed the expressional heterogeneity of cancer cells and stromal cells in MDA-MB-231 primary tumors and axillary lymph node metastases.

We observed two types of CSC-like populations in both the primary tumors and lymph node metastases. One of the CSC-like populations expressed CD44 and MYC. The CD44 gene is a well-known CSC marker in breast cancer [20, 37, 38]. The other CSC-like population, the HMGA1-high cluster, was observed in both the primary tumors and lymph node metastases. HMGA1 promotes tumor initiation, cancer stemness and metastasis in TNBC [19, 39, 40]. Enrichment analysis of DEGs showed upregulation of terms related to ribosome processes. The upregulation of translation and ribosomal processes may promote distant metastasis in breast cancer [41]. scRNA-seq analysis and Immunostaining of MDA-MB-231 xenografts confirmed the two types of CSC-like populations. In the integrated clinical scRNA-seq analysis, both types of CSC-like populations were observed in single-cell analysis of TNBC patients. Our results suggested that the coexistence of these multi-CSC-like populations makes curative treatment difficult and causes anticancer drug resistance in the clinic.

The marker genes and gene signatures identified from scRNA-seq studies have the potential to be prognosis markers in cancer patients [42]. The HMGA1 signatures with microspot resolution have the potential to be novel biomarkers for diagnosis, and HMGA1-high CSCs may contribute to poor prognosis. On the other hand, the CD44/MYC signatures did not correlate with progression. Previous research has reported that diagnosing invasive breast cancer on the basis of CD44 expression alone is difficult and that it is necessary to examine the expression of other genes [43, 44]. Our study supports that CD44/MYC signatures are not sufficient for diagnosis. In contrast, HMGA1 signatures are useful for the diagnosis of TNBC patients. The punching microdissection system used in this study enables the extraction of only the parts of a section, which is useful for the diagnosis of cancer and for analysis at low cost. This system could be applied in diagnosis instead of laser-capture microdissection methods.

Our results showed Inhba-high stromal populations in the xenograft model. Inhba is a member of the TGF-beta superfamily [45]. Inhba is upregulated in breast tumors and induces epithelial to mesenchymal transition (EMT), tumor growth and distant metastasis [46, 47]. Most Inhba-high populations also existed in the primary tumor. Our results suggest that stromal expression of Inhba enhanced tumor growth in the MDA-MB-231 primary tumors.

CCI analysis of each spot was performed to understand the interaction dynamics of different combinations of cancer cell and stromal cell types. B2M is a gene that presents self-antigens on the plasma membrane. Cancer cells present self-antigens to immune cells in the tumor tissue [48]. B2M has a different molecular regulatory mechanism in ER-positive and ER-negative breast cancer, and it controls the proliferation of cancer cells [49]. Our results suggest that MDA-MB-231 cells interact with immune cells through B2M antigen presentation, which controls cancer cell proliferation in the xenograft model.

Our study has limitations. Several spots did not contain enough RNA for analysis or exhibited bias toward either human RNA or mouse RNA (Supplementary Figure S1A). Thus, one limitation of this sampling method is that some spots have a biased cell type or no cells. Regarding the read counts (nCount_RNA) of each cluster, there were lower counts for the human Marker-low cluster, mouse Ttr-high cluster, mouse Inhba-high cluster, and mouse Bcell-like cluster than for other clusters (Supplementary Figure S1B and S1C). The limitations of cell type bias and low RNA extraction efficiency caused these low transcript counts. In particular, the marker-low human cluster and Ttr-high mouse cluster had the lowest nCount_RNA. This result suggests that these clusters may have included dead/dying cells. Tumor tissue has spatial heterogeneity, including in necroptotic/apoptotic areas, which is the one of the limitations of spatial transcriptome analysis. We also detected different nCount_RNA and features, so it was difficult to define the exact characteristics and cell types in the tissue at single-cell resolution. This is a technical limitation of spatial transcriptomics. More high-resolution spatial transcriptomics analyses are needed to define the cluster characteristics and cell–cell interactions at single-cell resolution, as has been recently developed [50].

In conclusion, our study showed that there are two types of CSC-like populations in MDA-MB-231 xenograft models and TNBC patients. The presence of these CSC-like populations could potentially make tumors more drug resistant, and more difficult to treat. More effective therapies need to be developed through the elucidation of intratumor heterogeneity. In addition, these spatial transcriptomics methods will be helpful for the diagnosis, further identification of biomarkers, and elucidation of the essential characteristics of cancer.

## Materials and Methods

### Cell culture

The MDA-MB-231-*luc2*-*Venus* cell line was cultured in RPMI-1640 medium (Fujifilm Wako, Osaka, Japan) supplemented with 10% heat-inactivated fetal bovine serum (FBS, Fujifilm Wako), 100 μg/ml streptomycin (Meiji Seika Pharma Co. Ltd. Tokyo, Japan) and 100 U/ml penicillin (Meiji Seika Pharma) at 37 °C with 5% CO_2_.

### Animal studies

A. breast cancer xenograft model was established in NOD.CB-17-Prkdc^scid^/J mice (NOD-SCID; Charles River Laboratories Japan, Inc., Kanagawa, Japan) by orthotopic transplantation as previously described [51]. A total of 1.0 x 10^6^ cells were injected into the 4th fat pad of NOD-SCID mice. The primary tumor was removed 8 weeks after transplantation. An axillary lymph node metastasis was sampled 2 weeks after removing the primary tumor. The growth of the primary tumors and metastases was monitored by bioluminescence using an in vivo imaging system (IVIS-XRMS, PerkinElmer, MA, USA). For bioluminescence monitoring by IVIS, mice were anesthetized with 2.5% isoflurane (Fujifilm Wako) and intraperitoneally injected with 3 mg D-luciferin (Gold Biotechnology Inc., MO, USA) in 200 μl PBS as previously described [52, 53]. The harvested organs were placed in ice-cold PBS (Fujifilm Wako), embedded in SCEM (Super Cryoembedding Medium, SECTION-LAB, Japan) using liquid nitrogen and stored at −80 °C until sectioning.

### Microtissue dissection and RNA-seq analysis

Microtissue sampling was performed by an automated tissue microdissection punching system as previously described [15]. Frozen sections were sliced at a thickness of 20 μm and transferred to an LMD film II (SECTIOIN-LAB). Microspots were sampled with a 100 μm needle in the dissection instrument. RNA-seq was performed by Illumina HiSeq as previously described [15].

### Mapping and quality check

Transcriptome analysis was performed with HISAT2 version 2.0.5 [16] and RSEM version 1.3.0 [54]. The gene expression of cancer cells was obtained by mapping RNA-sequence reads to the human reference genome or mouse reference genome. We subjected ‘protein_coding’ genes to spatial transcriptome analysis.

### Clustering and UMAP visualization

Data mining analyses such as clustering, UMAP analysis, and DEG extraction, were performed with the functions ‘runPCA’, ‘FindNeigbors’, ‘FindClusters’, and ‘runUMAP’ and ‘FindAllMarkers’ in ‘Seurat’ version 3.2. [18]. Cell cycle estimation was performed by the function ‘CellCycleScoring’ using cell cycle marker genes in Seurat. A heatmap of DEGs (adjusted p value < 0.05 and pct.1 - pct.2 > 0.1) (Supplementary Tables S2 and S4) was drawn using ‘ComplexHeatmap’ [55]. These packages and functions were run in R version 3.6.3.

### Network analysis for expressional heterogeneity

Correlational network analysis for calculation of the expressional heterogeneity was performed by the ‘igraph’ package as previously described [29, 30, 51]. We calculated Pearson’s correlational coefficients between spots classified into the same clusters. Next, we calculated the normalized closeness centrality using the correlational network.

### Enrichment analysis using DEGs

Pathway and GO enrichment analyses were performed by Metascape (https://metascape.org/gp/index.html#/main/step1) [21]. DEGs from each cluster were subjected to the Metascape interface. Differential enrichment terms were analyzed in multiple gene list mode. The results of the enrichment analysis were visualized as heatmaps.

### Immunostaining of tumor sections

Primary tumors generated from transplanted MDA-MB-231 cells were fixed in 4% paraformaldehyde two overnight and embedded in paraffin. Primary tumor sections were dewaxed with xylene and rehydrated with ethanol (100–70%). Antigen retrieval was performed by boiling the specimens in Immunosaver (Nissin EM, Tokyo, Japan) diluted 1:200 for 45 minutes at 98 °C. The sections were permeabilized with 0.1% Triton X-100 (Sigma-Aldrich Co. llc., MO, USA) for 15 minutes. After blocking with Dako blocking reagent (Vector Labs, CA, USA) for 30 minutes, sections were incubated with primary antibodies for 1 hour at room temperature. Sections were then incubated with the primary antibodies, anti-CD44 (60224-1-IG, Proteintech Group Inc., IL, USA, diluted 1:100) and anti-HMGA1 (AB252930, Abcam, Cambridge, UK, diluted 1:200) and secondary anti-mouse Alexa Fluor 488-labelled antibodies (Thermo Fisher Scientific, MA, USA, diluted 1:1000) and anti-rabbit Alexa Fluor 594-labelled antibodies (Thermo Fisher Scientific, diluted 1:1000). Slides were mounted with VECTASHIELD mounting medium with Hoechst 33342 (Thermo Fisher Scientific). Stained sections were imaged using an FV10i Laser Scanning Microscope (OLYMPUS, Tokyo, Japan).

### Analysis of the public scRNA-seq dataset

The scRNA-seq dataset of the MDA-MB-231-LM2 xenograft model (GSE163210) [28] was downloaded from the Gene Expression Omnibus (GEO) database. The scRNA-seq datasets of TNBC patients from the public datasets GSE161529 [31], GSE176078 [8], and GSE180286 [32] were also downloaded from the GEO database. The scRNA-seq datasets were analyzed with Seurat vignettes, and the integration of datasets was performed by Harmony [56]. Low-quality single cells (nFeature_RNA > 500 and percent.mt < 20) were removed. The clinical datasets were analyzed with clinical annotation data.

### Survival analysis

Survival analysis of the Molecular Taxonomy of Breast Cancer International Consortium (METABRIC) cohort [33, 57] was performed by the Kaplan–Meier method using the ‘ggplot2’, ‘survminer’ and ‘survival’ packages with R as previously described [58, 59].

### Cell–Cell Interaction (CCI) analysis

Ligand–receptor interactions between human cancer cells and mouse stromal cells were assessed using the interaction database of the Bader laboratory from Toronto University (https://baderlab.org/CellCellInteractions#Download_Data) in R. For spot CCI analysis, we extracted the genes whose expression value was greater than 2. We selected the combinations representing ligand–receptor interactions, in which both ligand genes and receptor genes were expressed in the same spot. Hierarchical clustering was performed by ‘hclust’, and the results were visualized as a circular clustering plot by the ‘circlize’ package in R [60].

### Statistical analysis

The chi-test, Fisher’s exact test, and the log rank test were performed in R. DEG extraction was performed by the ‘FindAllMarkers’ function with the Wilcoxon rank sum test. Significance was defined as P < 0.05.

### Code availability

The source code used for the spatial transcriptome analysis is available on GitHub (https://github.com/JunNakayama/Spatial-Transcriptomics-of-MDA-MB-231-xenografts)

### Data availability

The spatial expression data (RNA-seq data of each microspot) were deposited at GEO under accession number GSE184720.

## Acknowledgements

We thank Dr. Tomofumi Yamamoto, Mr. Kiyofumi Takahashi and Ms. Chikako Sakanashi for technical support. We thank Prof. Semba’s laboratory, Prof. Takeyama’s laboratory and CBBD-OIL (AIST) for meaningful discussion

## Authors’ Contributions

JN designed the study, performed the animal experiments and analysis of spatial transcriptomics, and wrote the manuscript. HM and TY performed the dissection of micro-tissues. KA performed analysis of NGS data. MH designed the experiment of micro-tissue dissection and interpreted the results. HT, YY and KS interpreted the results. All authors reviewed the manuscript.

## Ethics

Animal experiments in this study were conducted under the approval of the Animal Committee of Waseda University (2017-A043a).

## Supplementary Figure Legends

**Supplementary Figure S1.**
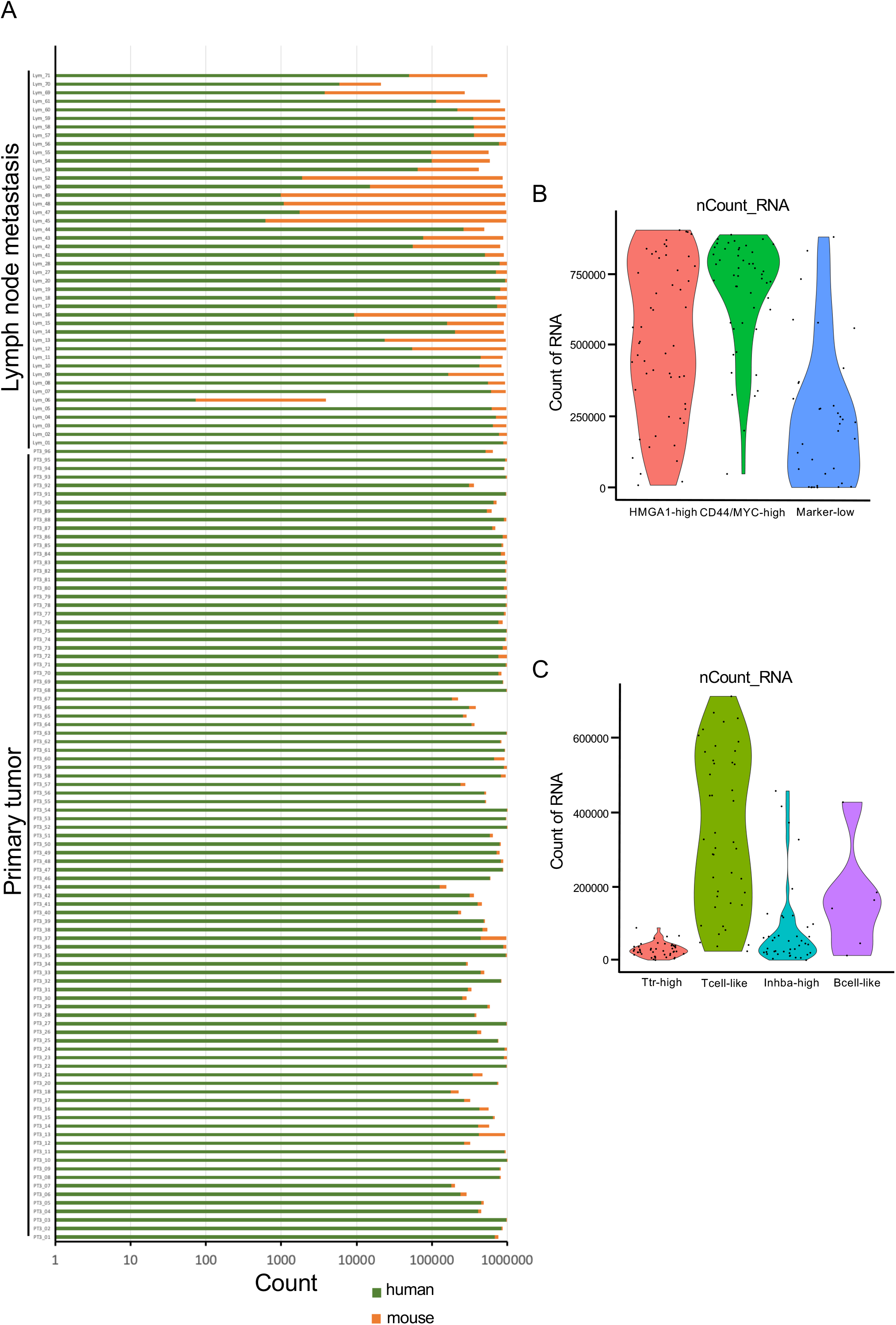
Read counts and heterogeneity in each spot. (A) Read counts of human and mouse transcripts in each spot. (B) Read counts of human cancer clusters. (C) Read counts of mouse stromal clusters.

**Supplementary Figure S2.**
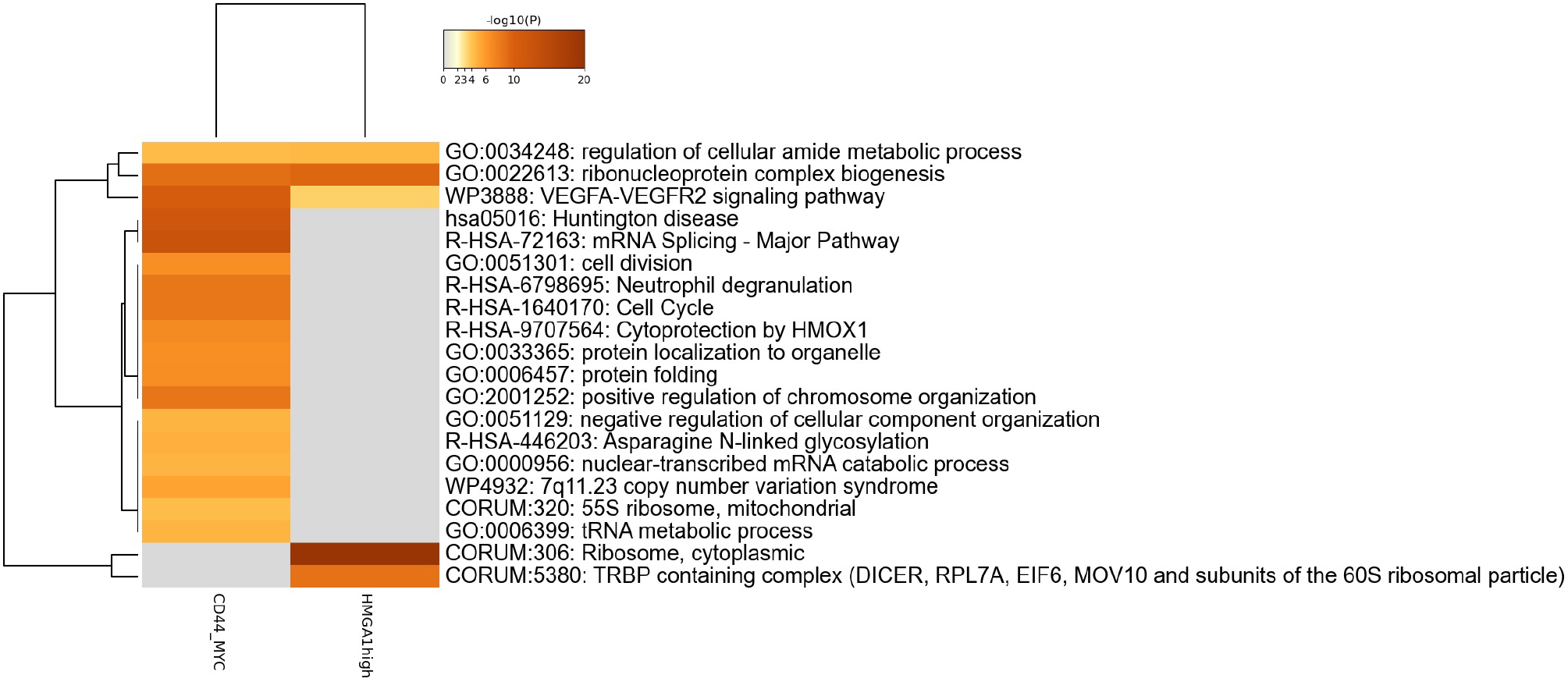
Enrichment analysis of DEGs in the two CSC-like populations. Heatmap of the results of the enrichment analysis of DEGs in the HMGA1-high and CD44/MYC-high clusters using Metascape.

**Supplementary Figure S3.**
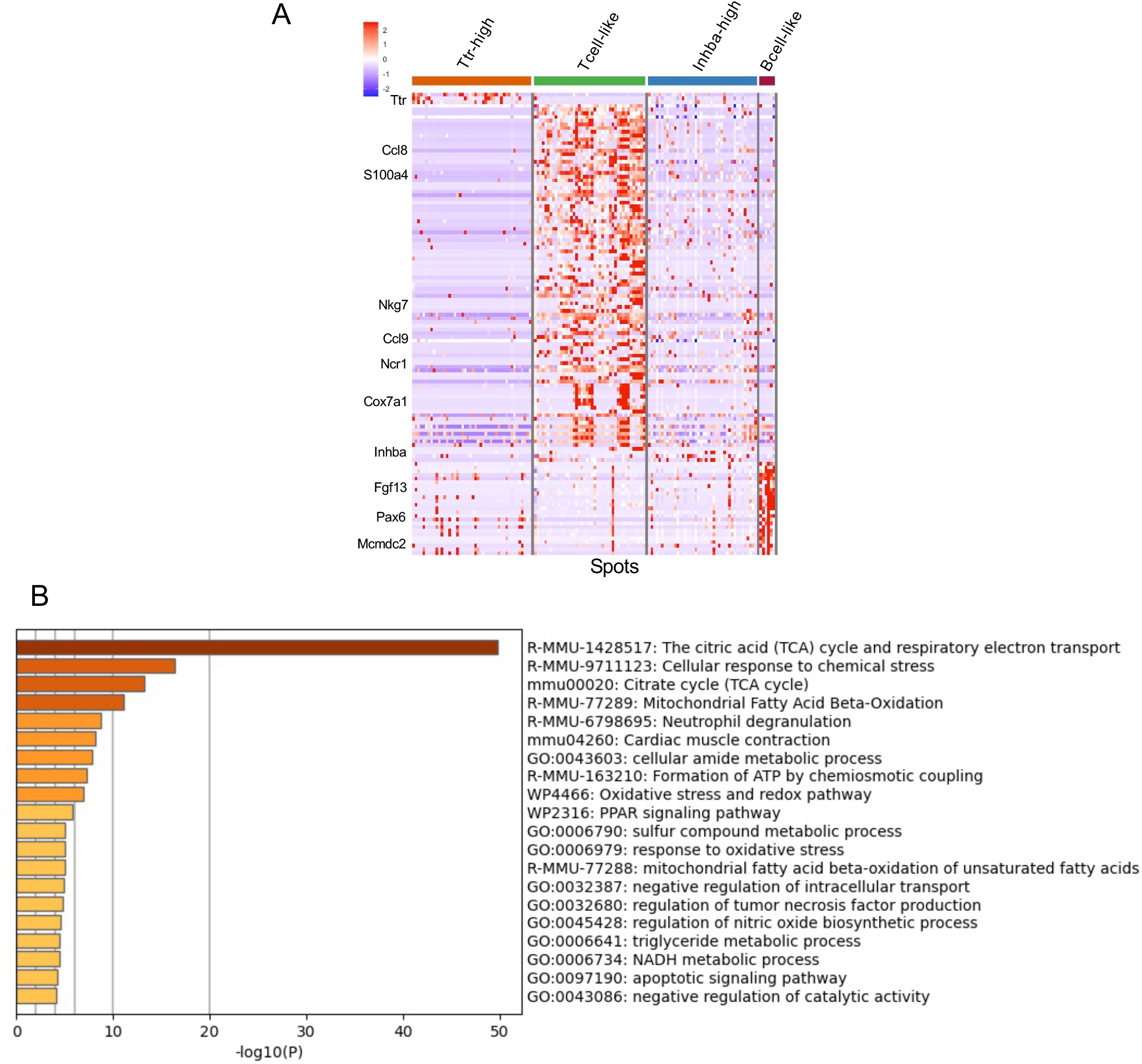
Stromal cluster DEG analysis and enrichment analysis. (A) Heatmap of the DEGs (adjusted p value < 0.05 and pct.1 - pct.2 > 0.1) in mouse stromal cells. (B) Heatmap of the results of the enrichment analysis in the Tcell-like cluster.

**Supplementary Figure S4.**
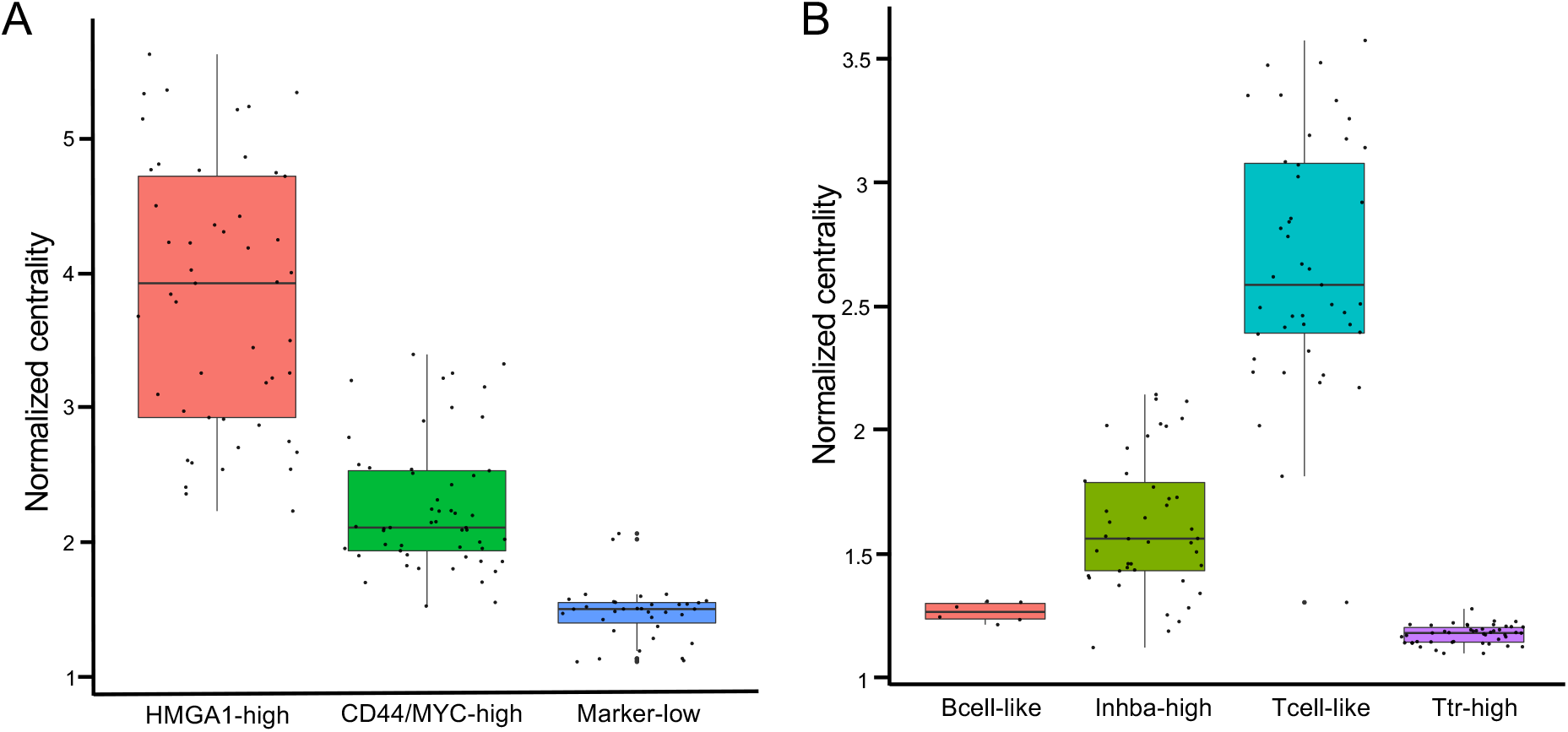
Closeness centrality for expressional heterogeneity analysis in each cluster. (A) Normalized closeness centrality in each human cancer cluster. (B) Normalized closeness centrality in each mouse stromal cluster.

**Supplementary Figure S5.**
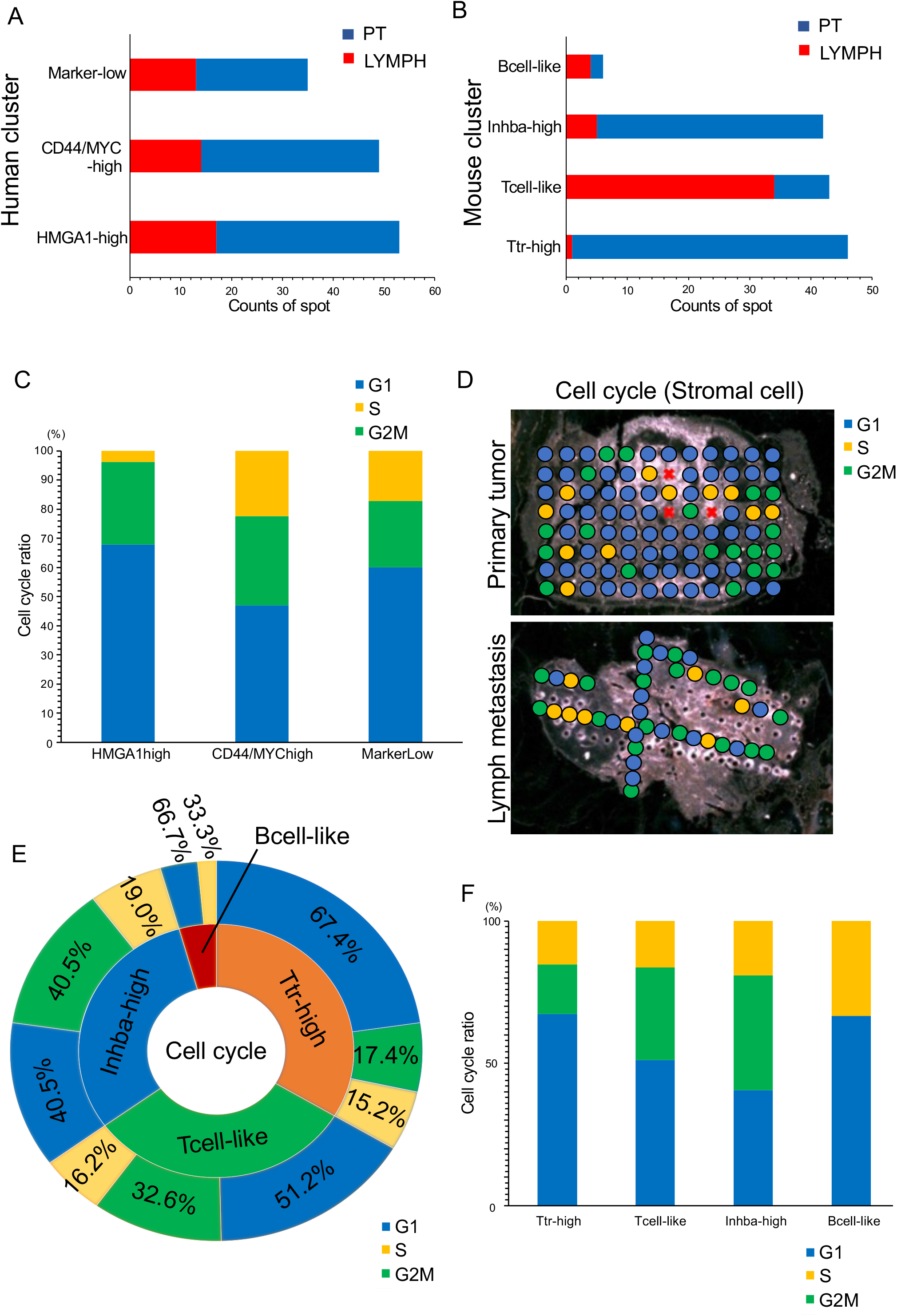
Spatial analysis of mouse clusters. (A and B) Bar plots of spot counts in the primary tumor and lymph node metastases: (A) human and (B) mouse. (C) Cell cycle phase proportions of the human cancer clusters. (D) Cell cycle proportions of the mouse stromal cells in the primary tumor and lymph node metastasis. (E) Sunburst plot of the cell cycle in mouse stromal cell clusters. (F) Cell cycle proportions of the mouse stromal clusters.

**Supplementary Figure S6.**
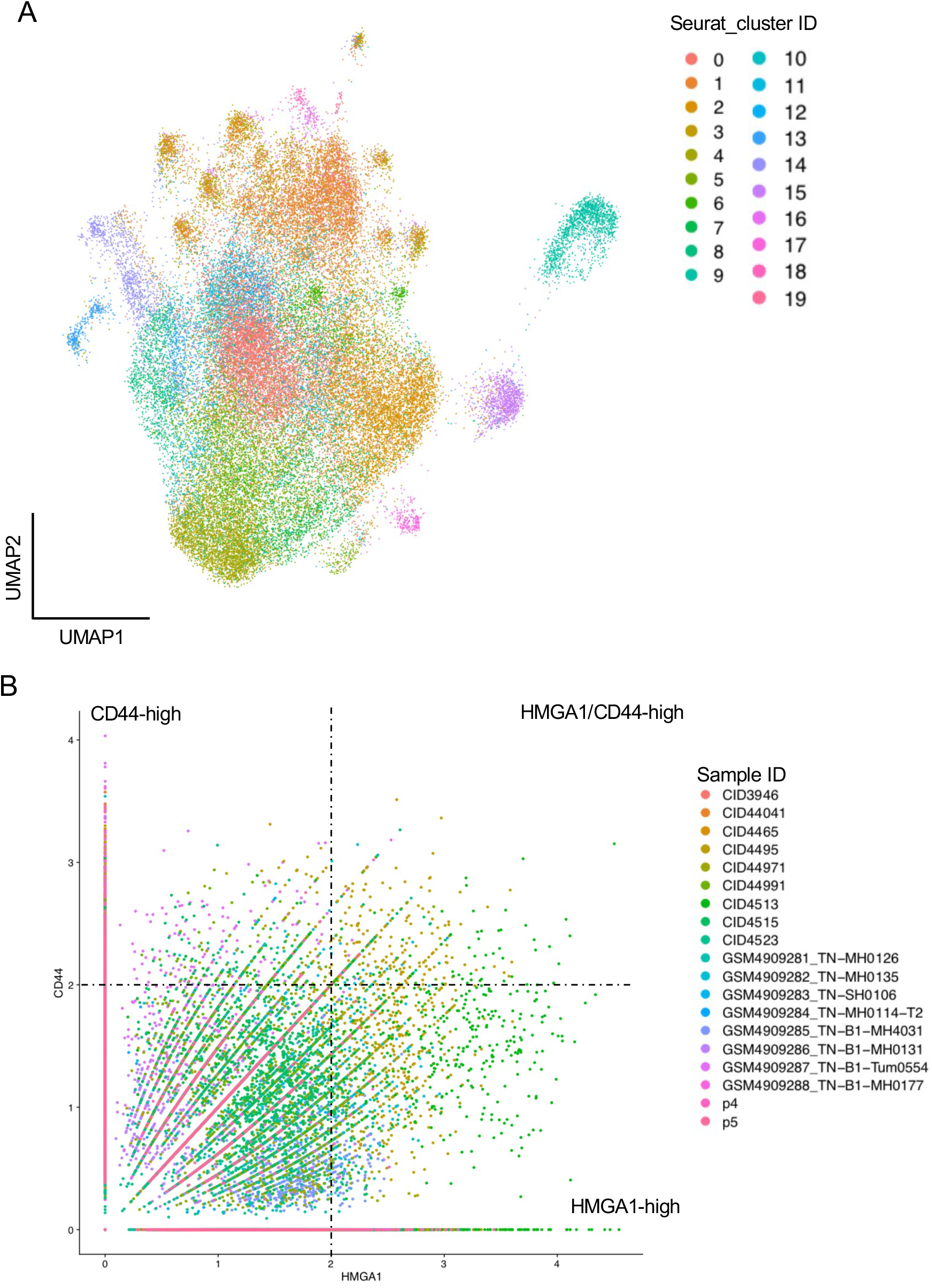
Reanalysis of integrated scRNA-seq data. (A) UMAP plot of cancer cells with cluster identification by Seurat. (B) Scatter plot of CD44 and HMGA1 expression in the integrated datasets.

**Supplementary Figure S7.**
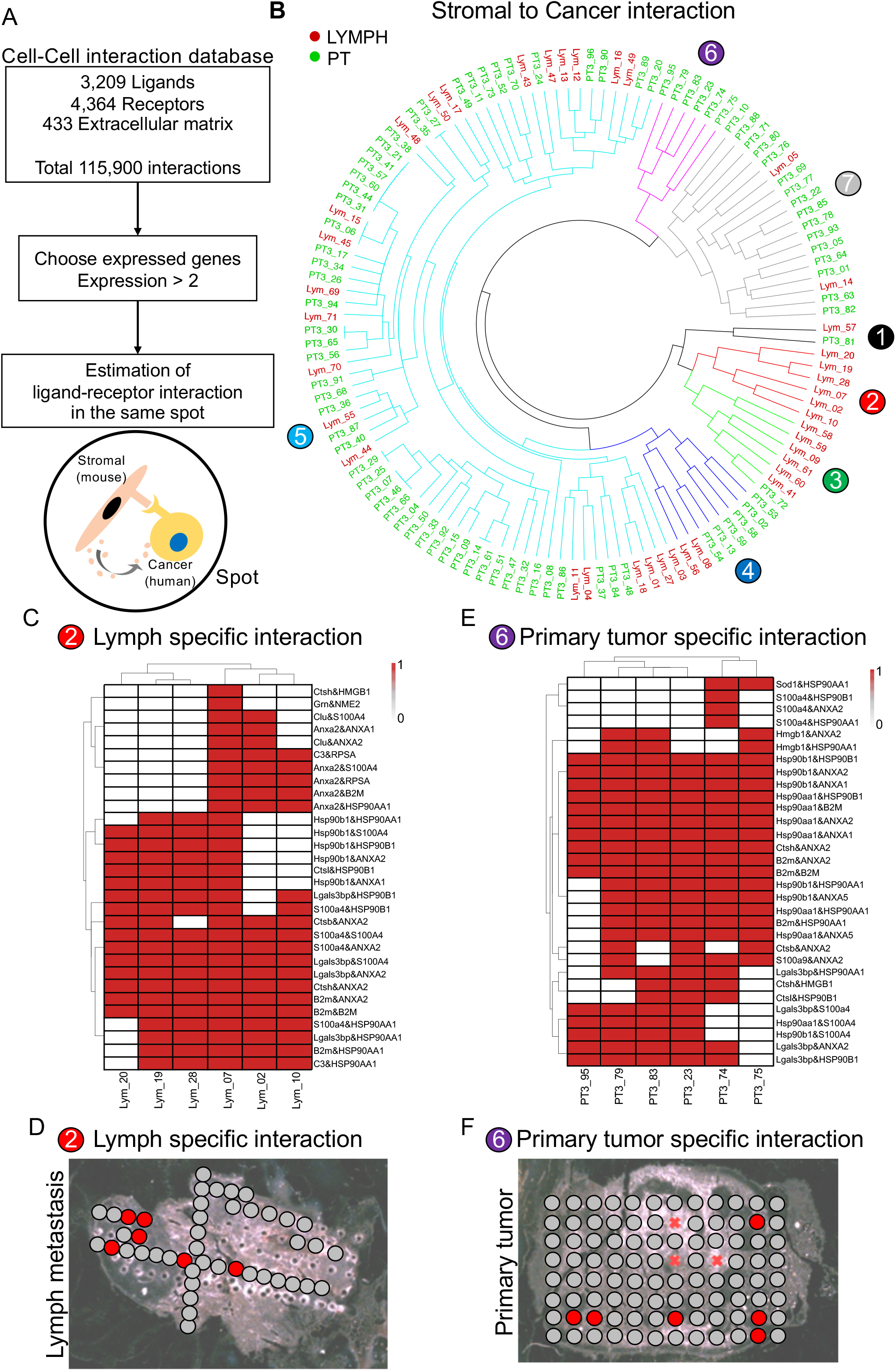
Spot Cell–Cell Interaction analysis of ‘cancer to stromal interactions’. (A) Flowchart of the CCI analysis. (B) Circular clustering plot of hierarchical analysis of spot CCI analysis (stromal cell to cancer cell). (C & E) Heatmap of CCI in the spots. 1: Detected interaction. 0: Not detected interaction. (D & F) Representative pictures show the position of CCI clusters.

**Supplementary Figure S8.**
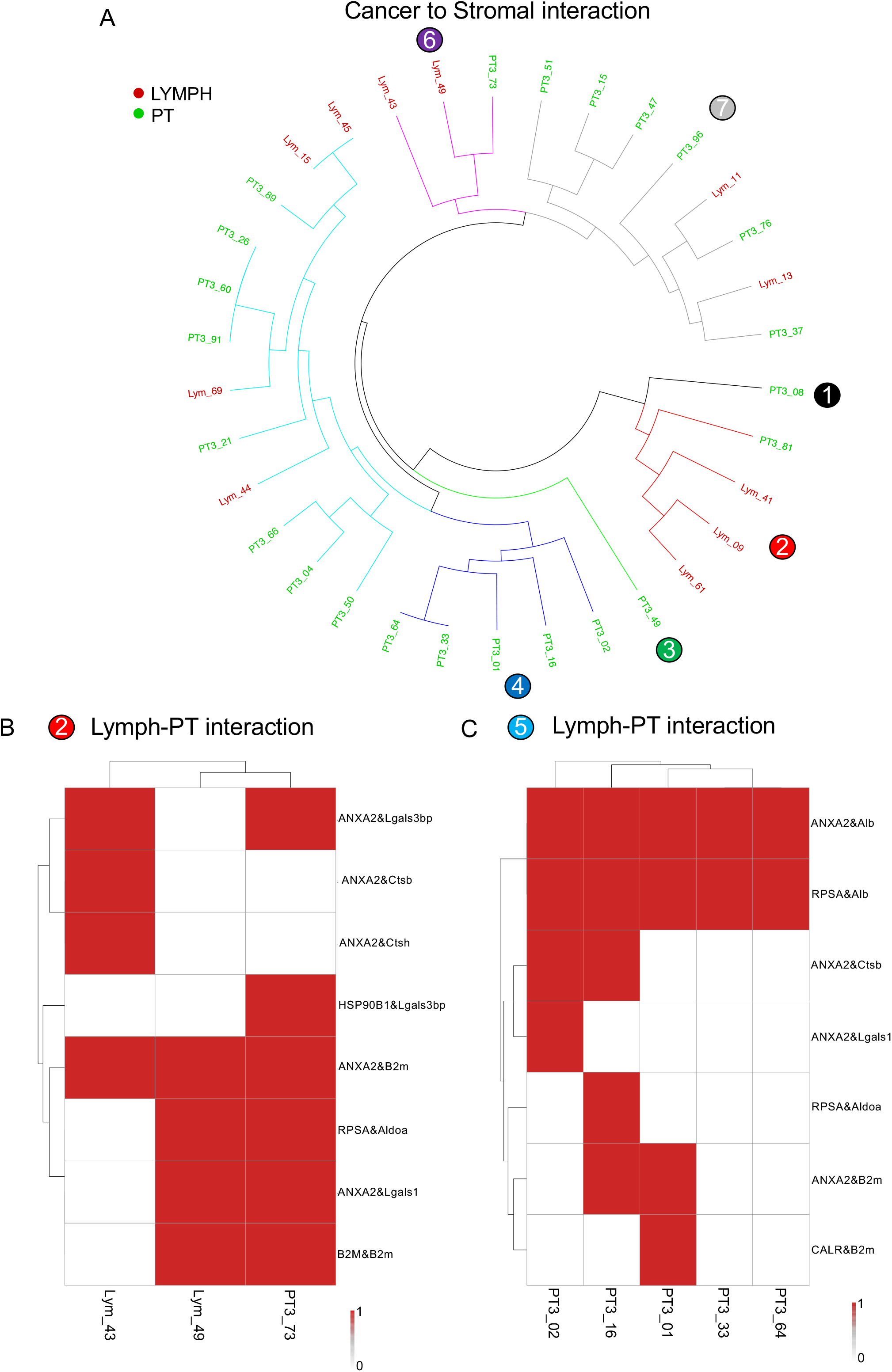
Spot Cell–Cell Interaction analysis of ‘cancer to stromal interactions’. (A) Circular clustering plot of hierarchical analysis of spot CCI analysis (cancer-to-stromal cell). (B & C) Heatmap of CCI in the interactive spots.

## Supplementary Tables

**Supplementary Table S1.** Clustering of microspots

**Supplementary Table S2.** Marker genes in human cancer cells

**Supplementary Table S3.** Upstream enrichment analysis of cancer cell clusters

**Supplementary Table S4.** Marker genes in mouse stromal cells

**Supplementary Table S5.** Cell cycle index in clusters

**Supplementary Table S6.** Cell annotation in TNBC patients

**Supplementary Table S7.** Signature gene lists

**Supplementary Table S8.** CCI list in clusters

## Reference

1. Obenauf AC, Massague J. Surviving at a Distance: Organ-Specific Metastasis. Trends Cancer. 2015;1(1):76–91. Epub 2015/09/01. doi: 10.1016/j.trecan.2015.07.009. PubMed PMID: 28741564; PubMed Central PMCID: PMC4673677.

2. Nakayama J, Han Y, Kuroiwa Y, Azuma K, Yamamoto Y, Semba K. The In Vivo Selection Method in Breast Cancer Metastasis. Int J Mol Sci. 2021;22(4). Epub 2021/03/07. doi: 10.3390/ijms22041886. PubMed PMID: 33672831; PubMed Central PMCID: PMC7918415.

3. Giuliano AE, Hunt KK, Ballman KV, Beitsch PD, Whitworth PW, Blumencranz PW, et al. Axillary dissection vs no axillary dissection in women with invasive breast cancer and sentinel node metastasis: a randomized clinical trial. Jama. 2011;305(6):569–75. Epub 2011/02/10. doi: 10.1001/jama.2011.90. PubMed PMID: 21304082; PubMed Central PMCID: PMC5389857.

4. McGranahan N, Swanton C. Clonal Heterogeneity and Tumor Evolution: Past, Present, and the Future. Cell. 2017;168(4):613–28. Epub 2017/02/12. doi: 10.1016/j.cell.2017.01.018. PubMed PMID: 28187284.

5. Yates LR, Knappskog S, Wedge D, Farmery JHR, Gonzalez S, Martincorena I, et al. Genomic Evolution of Breast Cancer Metastasis and Relapse. Cancer Cell. 2017;32(2):169–84. Epub 2017/08/16. doi: 10.1016/j.ccell.2017.07.005. PubMed PMID: 28810143; PubMed Central PMCID: PMC5559645.

6. Echeverria GV, Powell E, Seth S, Ge Z, Carugo A, Bristow C, et al. High-resolution clonal mapping of multi-organ metastasis in triple negative breast cancer. Nat Commun. 2018;9(1):5079. Epub 2018/12/01. doi: 10.1038/s41467-018-07406-4. PubMed PMID: 30498242; PubMed Central PMCID: PMC6265294.

7. Andersson A, Larsson L, Stenbeck L, Salmén F, Ehinger A, Wu SZ, et al. Spatial deconvolution of HER2-positive breast cancer delineates tumor-associated cell type interactions. Nat Commun. 2021;12(1):6012. Epub 2021/10/16. doi: 10.1038/s41467-021-26271-2. PubMed PMID: 34650042.

8. Wu SZ, Al-Eryani G, Roden DL, Junankar S, Harvey K, Andersson A, et al. A single-cell and spatially resolved atlas of human breast cancers. Nat Genet. 2021;53(9):1334–47. Epub 2021/09/09. doi: 10.1038/s41588-021-00911-1. PubMed PMID: 34493872.

9. Rao A, Barkley D, França GS, Yanai I. Exploring tissue architecture using spatial transcriptomics. Nature. 2021;596(7871):211–20. doi: 10.1038/s41586-021-03634-9.

10. Robinson DR, Wu YM, Lonigro RJ, Vats P, Cobain E, Everett J, et al. Integrative clinical genomics of metastatic cancer. Nature. 2017;548(7667):297–303. Epub 2017/08/08. doi: 10.1038/nature23306. PubMed PMID: 28783718; PubMed Central PMCID: PMC5995337.

11. Turdo A, Veschi V, Gaggianesi M, Chinnici A, Bianca P, Todaro M, et al. Meeting the Challenge of Targeting Cancer Stem Cells. Front Cell Dev Biol. 2019;7:16. Epub 2019/03/06. doi: 10.3389/fcell.2019.00016. PubMed PMID: 30834247; PubMed Central PMCID: PMC6387961.

12. Weiss F, Lauffenburger D, Friedl P. Towards targeting of shared mechanisms of cancer metastasis and therapy resistance. Nat Rev Cancer. 2022. Epub 2022/01/12. doi: 10.1038/s41568-021-00427-0. PubMed PMID: 35013601.

13. Oskarsson T, Batlle E, Massagué J. Metastatic Stem Cells: Sources, Niches, and Vital Pathways. Cell Stem Cell. 2014;14(3):306–21. doi: https://doi.org/10.1016/j.stem.2014.02.002.

14. Civita P, Franceschi S, Aretini P, Ortenzi V, Menicagli M, Lessi F, et al. Laser Capture Microdissection and RNA-Seq Analysis: High Sensitivity Approaches to Explain Histopathological Heterogeneity in Human Glioblastoma FFPE Archived Tissues. Front Oncol. 2019;9:482. Epub 2019/06/25. doi: 10.3389/fonc.2019.00482. PubMed PMID: 31231613; PubMed Central PMCID: PMC6568189.

15. Yoda T, Hosokawa M, Takahashi K, Sakanashi C, Takeyama H, Kambara H. Site-specific gene expression analysis using an automated tissue micro-dissection punching system. Sci Rep. 2017;7(1):4325. Epub 2017/07/01. doi: 10.1038/s41598-017-04616-6. PubMed PMID: 28659603; PubMed Central PMCID: PMC5489509.

16. Kim D, Paggi JM, Park C, Bennett C, Salzberg SL. Graph-based genome alignment and genotyping with HISAT2 and HISAT-genotype. Nat Biotechnol. 2019;37(8):907–15. Epub 2019/08/04. doi: 10.1038/s41587-019-0201-4. PubMed PMID: 31375807; PubMed Central PMCID: PMC7605509.

17. Butler A, Hoffman P, Smibert P, Papalexi E, Satija R. Integrating single-cell transcriptomic data across different conditions, technologies, and species. Nat Biotechnol. 2018;36(5):411–20. Epub 2018/04/03. doi: 10.1038/nbt.4096. PubMed PMID: 29608179; PubMed Central PMCID: PMC6700744.

18. Stuart T, Butler A, Hoffman P, Hafemeister C, Papalexi E, Mauck WM, 3rd, et al. Comprehensive Integration of Single-Cell Data. Cell. 2019; 177(7): 1888–902.e21. Epub 2019/06/11. doi: 10.1016/j.cell.2019.05.031. PubMed PMID: 31178118; PubMed Central PMCID: PMC6687398.

19. Pegoraro S, Ros G, Piazza S, Sommaggio R, Ciani Y, Rosato A, et al. HMGA1 promotes metastatic processes in basal-like breast cancer regulating EMT and stemness. Oncotarget. 2013;4(8): 1293–308. Epub 2013/08/16. doi: 10.18632/oncotarget.1136. PubMed PMID: 23945276; PubMed Central PMCID: PMC3787158.

20. Liu H, Patel MR, Prescher JA, Patsialou A, Qian D, Lin J, et al. Cancer stem cells from human breast tumors are involved in spontaneous metastases in orthotopic mouse models. Proc Natl Acad Sci U S A. 2010;107(42):18115–20. Epub 2010/10/06. doi: 10.1073/pnas.1006732107. PubMed PMID: 20921380; PubMed Central PMCID: PMC2964232.

21. Zhou Y, Zhou B, Pache L, Chang M, Khodabakhshi AH, Tanaseichuk O, et al. Metascape provides a biologist-oriented resource for the analysis of systems-level datasets. Nat Commun. 2019;10(1):1523. Epub 2019/04/05. doi: 10.1038/s41467-019-09234-6. PubMed PMID: 30944313; PubMed Central PMCID: PMC6447622.

22. Zhang X, Ren D, Guo L, Wang L, Wu S, Lin C, et al. Thymosin beta 10 is a key regulator of tumorigenesis and metastasis and a novel serum marker in breast cancer. Breast Cancer Res. 2017;19(1):15. Epub 2017/02/10. doi: 10.1186/s13058-016-0785-2. PubMed PMID: 28179017; PubMed Central PMCID: PMC5299657.

23. Ashraf Y, Mansouri H, Laurent-Matha V, Alcaraz LB, Roger P, Guiu S, et al. Immunotherapy of triple-negative breast cancer with cathepsin D-targeting antibodies. J Immunother Cancer. 2019;7(1):29. Epub 2019/02/06. doi: 10.1186/s40425-019-0498-z. PubMed PMID: 30717773; PubMed Central PMCID: PMC6360707.

24. Jung EJ, Moon HG, Cho BI, Jeong CY, Joo YT, Lee YJ, et al. Galectin-1 expression in cancer-associated stromal cells correlates tumor invasiveness and tumor progression in breast cancer. Int J Cancer. 2007;120(11):2331–8. Epub 2007/02/17. doi: 10.1002/ijc.22434. PubMed PMID: 17304502.

25. Balestrieri K, Kew K, McDaniel M, Ramez M, Pittman HK, Murray G, et al. Proteomic identification of tumor-and metastasis-associated galectin-1 in claudin-low breast cancer. Biochim Biophys Acta Gen Subj. 2021;1865(2):129784. Epub 2020/11/10. doi: 10.1016/j.bbagen.2020.129784. PubMed PMID: 33166603.

26. Lee YC, Huang CC, Lin DY, Chang WC, Lee KH. Overexpression of centromere protein K (CENPK) in ovarian cancer is correlated with poor patient survival and associated with predictive and prognostic relevance. PeerJ. 2015;3:e1386. Epub 2015/11/21. doi: 10.7717/peerj.1386. PubMed PMID: 26587348; PubMed Central PMCID: PMC4647587.

27. Wu H, Zhou Y, Wu H, Xu L, Yan Y, Tong X, et al. CENPN Acts as a Novel Biomarker that Correlates With the Malignant Phenotypes of Glioma Cells. Frontiers in Genetics. 2021;12. doi: 10.3389/fgene.2021.732376.

28. Moravec JC, Lanfear R, Spector DL, Diermeier SD, Gavryushkin A. Cancer phylogenetics using single-cell RNA-seq data. bioRxiv. 2021:2021.01.07.425804. doi: 10.1101/2021.01.07.425804.

29. Watanabe N, Nakayama J, Fujita Y, Mori Y, Kadota T, Shimomura I, et al. Single-cell Transcriptome Analysis Reveals an Anomalous Epithelial Variation and Ectopic Inflammatory Response in Chronic Obstructive Pulmonary Disease. medRxiv. 2020:2020.12.03.20242412. doi: 10.1101/2020.12.03.20242412.

30. Nakayama J, Yamamoto Y. Single-cell meta-analysis of cigarette smoking lung atlas. bioRxiv. 2021:2021.12.09.472029. doi: 10.1101/2021.12.09.472029.

31. Pal B, Chen Y, Vaillant F, Capaldo BD, Joyce R, Song X, et al. A single-cell RNA expression atlas of normal, preneoplastic and tumorigenic states in the human breast. Embo j. 2021;40(11):e107333. Epub 2021/05/06. doi: 10.15252/embj.2020107333. PubMed PMID: 33950524; PubMed Central PMCID: PMC8167363.

32. Xu K, Wang R, Xie H, Hu L, Wang C, Xu J, et al. Single-cell RNA sequencing reveals cell heterogeneity and transcriptome profile of breast cancer lymph node metastasis. Oncogenesis. 2021;10(10):66. Epub 2021/10/07. doi: 10.1038/s41389-021-00355-6. PubMed PMID: 34611125; PubMed Central PMCID: PMC8492772.

33. Pereira B, Chin S-F, Rueda OM, Vollan H-KM, Provenzano E, Bardwell HA, et al. The somatic mutation profiles of 2,433 breast cancers refine their genomic and transcriptomic landscapes. Nat Commun. 2016;7(1):11479. doi: 10.1038/ncomms11479.

34. Steeg PS. Targeting metastasis. Nat Rev Cancer. 2016;16(4):201–18. Epub 2016/03/25. doi: 10.1038/nrc.2016.25. PubMed PMID: 27009393; PubMed Central PMCID: PMC7055530.

35. Bradford JR, Farren M, Powell SJ, Runswick S, Weston SL, Brown H, et al. RNA-Seq Differentiates Tumour and Host mRNA Expression Changes Induced by Treatment of Human Tumour Xenografts with the VEGFR Tyrosine Kinase Inhibitor Cediranib. PLOS ONE. 2013;8(6):e66003. doi: 10.1371/journal.pone.0066003.

36. Callari M, Batra AS, Batra RN, Sammut S-J, Greenwood W, Clifford H, et al. Computational approach to discriminate human and mouse sequences in patient-derived tumour xenografts. BMC Genomics. 2018;19(1):19-. doi: 10.1186/s12864-017-4414-y. PubMed PMID: 29304755.

37. Marotta LL, Almendro V, Marusyk A, Shipitsin M, Schemme J, Walker SR, et al. The JAK2/STAT3 signaling pathway is required for growth of CD44^+^CD24^-^ stem cell-like breast cancer cells in human tumors. J Clin Invest. 2011;121(7):2723–35. Epub 2011/06/03. doi: 10.1172/jci44745. PubMed PMID: 21633165; PubMed Central PMCID: PMC3223826.

38. Sheridan C, Kishimoto H, Fuchs RK, Mehrotra S, Bhat-Nakshatri P, Turner CH, et al. CD44+/CD24-breast cancer cells exhibit enhanced invasive properties: an early step necessary for metastasis. Breast Cancer Res. 2006;8(5):R59. Epub 2006/10/26. doi: 10.1186/bcr1610. PubMed PMID: 17062128; PubMed Central PMCID: PMC1779499.

39. Huang R, Huang D, Dai W, Yang F. Overexpression of HMGA1 correlates with the malignant status and prognosis of breast cancer. Mol Cell Biochem. 2015;404(1-2):251–7. Epub 2015/03/17. doi: 10.1007/s11010-015-2384-4. PubMed PMID: 25772486.

40. Shah SN, Cope L, Poh W, Belton A, Roy S, Talbot CC, Jr., et al. HMGA1: a master regulator of tumor progression in triple-negative breast cancer cells. PLoS One. 2013;8(5):e63419. Epub 2013/05/10. doi: 10.1371/journal.pone.0063419. PubMed PMID: 23658826; PubMed Central PMCID: PMC3642138.

41. Ebright RY, Lee S, Wittner BS, Niederhoffer KL, Nicholson BT, Bardia A, et al. Deregulation of ribosomal protein expression and translation promotes breast cancer metastasis. Science. 2020;367(6485): 1468–73. Epub 2020/02/08. doi: 10.1126/science.aay0939. PubMed PMID: 32029688; PubMed Central PMCID: PMC7307008.

42. Cheng S, Li Z, Gao R, Xing B, Gao Y, Yang Y, et al. A pan-cancer single-cell transcriptional atlas of tumor infiltrating myeloid cells. Cell. 2021;184(3):792–809.e23. Epub 2021/02/06. doi: 10.1016/j.cell.2021.01.010. PubMed PMID: 33545035.

43. Wang H, Wang L, Song Y, Wang S, Huang X, Xuan Q, et al. CD44(+)/CD24(-) phenotype predicts a poor prognosis in triple-negative breast cancer. Oncol Lett. 2017;14(5):5890–8. Epub 2017/11/09. doi: 10.3892/ol.2017.6959. PubMed PMID: 29113223; PubMed Central PMCID: PMC5661458.

44. Mylona E, Giannopoulou I, Fasomytakis E, Nomikos A, Magkou C, Bakarakos P, et al. The clinicopathologic and prognostic significance of CD44+/CD24(-/low) and CD44-/CD24+ tumor cells in invasive breast carcinomas. Hum Pathol. 2008;39(7):1096–102. Epub 2008/05/23. doi: 10.1016/j.humpath.2007.12.003. PubMed PMID: 18495204.

45. Bloise E, Ciarmela P, Dela Cruz C, Luisi S, Petraglia F, Reis FM. Activin A in Mammalian Physiology. Physiol Rev. 2019;99(1):739–80. Epub 2018/12/13. doi: 10.1152/physrev.00002.2018. PubMed PMID: 30540228.

46. Bashir M, Damineni S, Mukherjee G, Kondaiah P. Activin-A signaling promotes epithelial-mesenchymal transition, invasion, and metastatic growth of breast cancer. NPJ Breast Cancer. 2015;1:15007. Epub 2015/08/12. doi: 10.1038/npjbcancer.2015.7. PubMed PMID: 28721365; PubMed Central PMCID: PMC5515205.

47. Kalli M, Mpekris F, Wong CK, Panagi M, Ozturk S, Thiagalingam S, et al. Activin A Signaling Regulates IL13Rα2 Expression to Promote Breast Cancer Metastasis. Front Oncol. 2019;9:32. Epub 2019/02/26. doi: 10.3389/fonc.2019.00032. PubMed PMID: 30805303; PubMed Central PMCID: PMC6370707.

48. Popat S, Grohé C, Corral J, Reck M, Novello S, Gottfried M, et al. Anti-angiogenic agents in the age of resistance to immune checkpoint inhibitors: Do they have a role in non-oncogene-addicted non-small cell lung cancer? Lung Cancer. 2020;144:76–84. Epub 2020/05/11. doi: 10.1016/j.lungcan.2020.04.009. PubMed PMID: 32387684.

49. Chai D, Li K, Du H, Yang S, Yang R, Xu Y, et al. β2-microglobulin has a different regulatory molecular mechanism between ER(+) and ER(-) breast cancer with HER2(). BMC Cancer. 2019;19(1):223. Epub 2019/03/15. doi: 10.1186/s12885-019-5410-1. PubMed PMID: 30866857; PubMed Central PMCID: PMC6417228.

50. Srivatsan SR, Regier MC, Barkan E, Franks JM, Packer JS, Grosjean P, et al. Embryo-scale, single-cell spatial transcriptomics. Science. 2021;373(6550):111–7. Epub 2021/07/03. doi: 10.1126/science.abb9536. PubMed PMID: 34210887.

51. Nakayama J, Ito E, Fujimoto J, Watanabe S, Semba K. Comparative analysis of gene regulatory networks of highly metastatic breast cancer cells established by orthotopic transplantation and intra-circulation injection. Int J Oncol. 2017;50(2):497–504. Epub 2016/12/22. doi: 10.3892/ijo.2016.3809. PubMed PMID: 28000849.

52. Han Y, Nakayama J, Hayashi Y, Jeong S, Futakuchi M, Ito E, et al. Establishment and characterization of highly osteolytic luminal breast cancer cell lines by intracaudal arterial injection. Genes Cells. 2020;25(2):111–23. Epub 2019/12/19. doi: 10.1111/gtc.12743. PubMed PMID: 31849141.

53. Kuroiwa Y, Nakayama J, Adachi C, Inoue T, Watanabe S, Semba K. Proliferative Classification of Intracranially Injected HER2-positive Breast Cancer Cell Lines. Cancers (Basel). 2020;12(7):1811. Epub 2020/07/10. doi: 10.3390/cancers12071811. PubMed PMID: 32640677; PubMed Central PMCID: PMC7408688.

54. Li B, Dewey CN. RSEM: accurate transcript quantification from RNA-Seq data with or without a reference genome. BMC Bioinformatics. 2011;12:323. Epub 2011/08/06. doi: 10.1186/1471-2105-12-323. PubMed PMID: 21816040; PubMed Central PMCID: PMC3163565.

55. Gu Z, Eils R, Schlesner M. Complex heatmaps reveal patterns and correlations in multidimensional genomic data. Bioinformatics. 2016;32(18):2847–9. doi: 10.1093/bioinformatics/btw313.

56. Korsunsky I, Millard N, Fan J, Slowikowski K, Zhang F, Wei K, et al. Fast, sensitive and accurate integration of single-cell data with Harmony. Nat Methods. 2019;16(12):1289–96. Epub 2019/11/20. doi: 10.1038/s41592-019-0619-0. PubMed PMID: 31740819; PubMed Central PMCID: PMC6884693.

57. Curtis C, Shah SP, Chin SF, Turashvili G, Rueda OM, Dunning MJ, et al. The genomic and transcriptomic architecture of 2,000 breast tumours reveals novel subgroups. Nature. 2012;486(7403):346–52. Epub 2012/04/24. doi: 10.1038/nature10983. PubMed PMID: 22522925; PubMed Central PMCID: PMC3440846.

58. Murakami A, Maekawa M, Kawai K, Nakayama J, Araki N, Semba K, et al. Cullin-3/KCTD10 E3 complex is essential for Rac1 activation through RhoB degradation in human epidermal growth factor receptor 2-positive breast cancer cells. Cancer Sci. 2019;110(2):650–61. Epub 2018/12/06. doi: 10.1111/cas.13899. PubMed PMID: 30515933; PubMed Central PMCID: PMC6361568.

59. Nishiyama K, Maekawa M, Nakagita T, Nakayama J, Kiyoi T, Chosei M, et al. CNKSR1 serves as a scaffold to activate an EGFR phosphatase via exclusive interaction with RhoB-GTP. Life Sci Alliance. 2021;4(9). Epub 2021/07/01. doi: 10.26508/lsa.202101095. PubMed PMID: 34187934; PubMed Central PMCID: PMC8321701.

60. Gu Z, Gu L, Eils R, Schlesner M, Brors B. circlize implements and enhances circular visualization in R. Bioinformatics. 2014;30(19):2811–2. doi: 10.1093/bioinformatics/btu393.

